# Influence of fats, carotenoids and POPs on the reproduction of the leatherback turtle

**DOI:** 10.1101/669580

**Authors:** E. De Andrés, Juan José Negro Balmaseda, Juan Carlos Navarro, Adolfo Marco

## Abstract

This study provides baseline data on fatty acid profiles and carotenoids in eggs of the leatherback sea turtle, *Dermochelys coriacea*. Correlations among such compounds, persistent organic pollutants, and reproductive parameters are also investigated. A number of 18 clutches were evaluated during June and August of 2008 at Reserva Pacuare Beach, in the Caribbean side of Costa Rica. Viability and fertility were calculated for each nest. Concentration levels of saturated and unsaturated fatty acids (SFAs and PUFAs), carotenoids and different congeners of persistent organic pollutants (POPs) were determined from egg-yolk samples. Mean ± SD values were calculated for each compound and for each clutch. Correlations were performed searching for interactions among different compounds and for potential effects on reproductive parameters, thus all the studied compounds were related to each other and to any of the reproductive parameters. Low carotenoid levels characterized all eggs of this species, and a positive relationship between carotenoid concentrations and the viability rate was found. POPs and PUFA were positive and strongly correlated, suggesting molecular interactions. PUFAs appeared to increase fertility rate and hatchling length. This study provides potential evidences of PUFA enhancing fertility and hatchling size, and of carotenoids limiting vitellogenesis. The positive correlation found between POPs and PUFAs may indicate that harmful effects of these contaminants on the reproduction of leatherback turtles could be masked.

## INTRODUCTION

The leatherback turtle (*Dermochelys coriacea*) is the largest of the sea turtle species, reaching 2 m in length, and the only of them capable to dive up to 1000 m depth [1]. During the past three decades, concern about this species has increased due to the diminishing trend on nesting females of the East Pacific subpopulations, which have decreased 97.4% during the past three generations [2]. That was the main reason to globally catalog it first as a Critically Endangered Species, and that Pacific subpopulation is thought to be no longer recovered [3. On the other hand, Caribbean nesting subpopulations of *D. coriacea* appear to be increasing by 20.6% over the past three generations [4], probably due to an intensive program of protection and egg relocation carried out for more than 20 years at certain locations [5]. Nevertheless, although Caribbean nesting population sizes are considered to be very small if compared to those that nested in the Pacific coasts less than ten years ago (when these Pacific populations were already endangered), it is currently considered as a vulnerable species regarding to the last published IUCN report [3]. The causes of the global decreasing trend in *D. coriacea* may be due to biological limitations inherent in the reproduction of the leatherback turtle, the effects derived from anthropogenic activities or a combination of both. Leatherback females reach the age of maturity at about 9 years old and nest on tropical and subtropical waters worldwide every two to three years [6]. They are able to lay up to 11 nests about ten days apart within a single nesting season with clutches ranging from 65 to 86 eggs [7, 8, 9]. Fertility rate in *Dermochelys coriacea* is usually high, reaching around 70-90%. Nevertheless, hatch success has been reported to be lower than those of other sea turtle species with values < 60% [10, 11, 12, 13]. Several threats have been identified to leatherback clutches during incubation such as beach erosion, high sand water content, high incubation temperature or beach pollution [14, 15, 16, 17] though those threats are also affecting to the rest of sea turtle species. The special feeding behavior and diet of the leatherback turtle may affect the egg yolk quality and subsequently could cause their low hatching success.

Although it is not clear whether they opportunistically eat or not during breeding season [18, 19], it is well known that leatherback turtle migrates long distances from nesting beaches towards foraging areas in order to supply reproduction requirements and keep fat reserves for the next nesting season [20, 21]. These areas where leatherbacks migrate to are located in northern and southern latitudes, where waters are colder and more productive. However, turtles are relatively slow and are not able to predate large pelagic vertebrates and their diet consists primarily on gelatinous zooplankton, focusing mainly in large species such as the lion’s mane jelly (*Cyanea capillata*) although they also eat smaller species [22, 23, 24]. During this feeding season, leatherback turtles have an intake of essential nutrients that will serve as a basis for the egg production. Egg yolk nutrient composition consists in proteins, fatty acids, fat-soluble antioxidants (vitamins A, E and carotenoids) and lipids in general. These compounds are strictly necessary and play an important role in embryonic development in turtles [25, 26, 27, 28]; even more important for this species since low reproductive success is due to high embryonic mortality [8]. However, there is a lack of knowledge in this field to date since no studies have been carried out to determine baseline levels of carotenoids or fatty acid profiles characterizing leatherback turtle eggs, neither their potential effects on reproductive success.

Currently, many of the hazards affecting leatherback turtles are linked to human activities such as fishery bycatch, illegal egg collection, plastic ingestion or contamination of foraging areas [29, 30]. Different studies about POP contamination evidenced that certain levels of POPs have negative effects on health parameters of sea turtles, that these contaminants are maternally transferred, that they may negatively interfere in the right embryonic development and hatchling sizes, and that they might reduce hatching success rates [31, 32, 33, 34, 35, 36]. Due to their lipophilic nature and the mobilization of lipids by sea turtles for egg production [37], these POPs may also interact or be related to other lipophilic compounds that are essential nutrients for the embryo development such as carotenoids or fatty acids. In fact, POP concentrations in egg yolk of *C. caretta* can be doubled by the late stages of embryonic development, consistently with the 50% increase in lipid content [32]. Thus, if those interactions occur, hatching success and hatchling survival could be compromised and population of leatherback turtles may tend to decrease even more. Therefore, in order to know at what point populations of leatherback turtles are at present, these potential relations and implications need to be studied. This study complements the work published in [36]. The primary goals here are: 1) to provide additional baseline data on lipid content, fatty acid profiles, and carotenoid composition in eggs of the leatherback turtle; 2) to investigate whether any associations among these different compounds and POPs exist; and 3) to clear up how any of these compounds may be related to viability, fertility, hatch success or hatchling morphometrics in this species.

## METHODOLOGY

### Sampling

The ethics committee of the National System of Protected Areas of the Ministry of Environment and Energy of Costa Rica prospectively approved this research in the resolution 009-2008-INV-ACLAC. A total of 18 nests of different leatherback turtles were evaluated during the period between June and August of 2008, along the beaches of Pacuare Reserve (Caribbean side of Costa Rica). This nesting area has a high density of leatherback nests within one of the largest leatherback rookeries of the world [38, 39]. Curved carapace length (CCL), curved carapace width (CCW), clutch size (number of potentially fertile eggs) and number of shelled albumin globes (SAGs) were determined at the time of oviposition [8], from 3^rd^ to 8^th^ of June. At the same time, one to three eggs were collected from each nest for chemical analysis. The eggs were sent to the Doñana Biological Station of Spain by plane in an isotherm container with the CITES permit 08CR000006/SJ. In early August emergence started. Hatching success, fertility and viability rates were calculated following De Andrés et al [36]. Immediately after emergence, hatchling mass (±0.1 gr), straight carapace length (SLC) and straight carapace width (SCW) were successfully measured (±0.1 mm) in only eight of the clutches.

### Chemical analysis

Total lipids from freeze dried egg samples, were extracted with chloroform:methanol (2:1, v/v), following the method described by Folch *et al*. [40]. After evaporation of the solvent under nitrogen, the lipid fractions were dried overnight in a vacuum desiccator, and quantified gravimetrically. Total lipids were dissolved in chloroform:methanol (2:1, v/v) with 0.01% of butylated hydroxytoluene (BHT) added as an antioxidant, and stored at −25 °C, in sealed vials, under nitrogen.

Fatty acid methyl esters (FAME) were prepared from the total lipid fractions by acid-catalyzed transmethylation [41]. Nonadecanoic acid (19:0) was used as an internal standard for quantification purposes. After extraction with hexane:diethyl ether (1:1, v/v), FAME were subsequently purified by thin layer chromatography (Silca gel G 60, Merck) using hexane:diethyl:ether:acetic acid (85:15:1.5, v/v/v) as the solvent phase. The purified methyl esters were then dissolved in hexane containing 0.01% BHT and analysed in a gas chromatography (Fisons Instruments 8000 series GC, Italy) equipped with a flame ionization detector (FID), a silica capillary column (30 m × 0.25 mm × 0.25 µm film thicknessTR-WAX, Teknokroma, Spain), and a cold on-column injection system, using helium as carrier gas. Samples were ran under thermal gradient from 50 to 220°C. The identification of the peaks was carried out by comparison with well characterised standards. Carotenoid extraction from egg yolk was performed [42]. Briefly, egg yolk (0.2–0.5 g) was homogenized in 2 mL of 1:1 (v/v) mixture of 5% NaCl solution and ethanol, followed by the addition of 3 mL hexane and further homogenization for 3 min. After centrifugation hexane was collected and the extraction was repeated twice. Hexane extracts were combined and evaporated under N_2_, and the residue was dissolved in 1 mL of methanol:dichloromethane (1:1, v/v) and centrifuged, and the supernatant was used for carotenoid determination. Carotenoids were determined by high performance liquid chromatography as described previously [43](Surai *et al*. 2000), using a Spherisorb type S3ODS2, 5-µm C^18^, reverse-phase column, 25 cm × #4.6 mm (Phase Separation, Clwyd, U.K.) with a mobile phase of acetonitrile:methanol (85:15) and acetonitrile:dichloromethane:methanol (70:20:10) in gradient elution using detection by absorbance at 445 nm. Peaks were identified by comparison with the retention times of a range of carotenoid standards (variously obtained from Sigma, Poole, U.K.; Fluka, Gillingham, U.K.; Apin, Abingdon, U.K.; and Hoffman-La Roche, Basel, Switzerland), as well as using coevolution of individual carotenoids with known standards.

We searched for different congeners of PCBs (polychlorinated biphenyls) and PBDEs (polybrominated diphenyl ethers) present in the eggs. Extraction, purification and determination techniques for quantification of the different POPs are thoroughly described in De Andrés et al [36].

### Data Analysis

For base-line data of the different compounds, all concentrations and proportions were averaged between all clutches. Mean ± standard deviation (SD), median, minimum and maximum values were calculated for all compound groups. Grubbs’ test was applied to searching for potential outliers. To evaluate those potential outliers, values were compared to the rest of eggs, to the eggs belonging to the same nest, to the patterns observed within the other nests and to other values reported in the literature. STATISTICA (v 8.0) was used to carry out statistical descriptions, normality tests, Grubbs’ test and correlations.

In order to evaluate differences among clutches and determine whether egg sampling was representative enough, analysis of similarity (ANOSIM) under Bray-Curtis index was performed, separately, for all lipids and carotenoids with no data transformation using PRIMER (v 6.1.6). Each class or compound was entered as a separated variable and each egg was analyzed as an individual sample. To better illustrate the differences between and within clutches, a non-metric multi-dimensional scaling (NMDS) plot was also constructed using PRIMER (v 6.1.6) [34, 36].

Arcsine-transformation was applied to all proportional data such as reproductive rates (viability, fertility and hatching success) as well as fatty acid proportions [8, 44]. Shapiro-Wilk Normality test was performed for all variables in order to choose the best possible analysis. To investigate any relationship between different compounds, parametric correlations were mainly performed, except for those variables that exhibited a non-normal distribution (Shapiro-Wilk test, *p* < 0.05), where non-parametric correlations were then executed. When possible, parametric correlations were also used to evaluate whether it exists or not any potential influence (relationship) of these compounds on reproductive parameters or hatchling morphometrics.

## RESULTS

### Reproductive parameters and POPs

Results are synthesized in Table 1. Values are averaged by nest and total mean calculated for each variable. Mean clutch size (i.e. number of yolked eggs) was 79 ±11 eggs, with viability and fertility rates being of 0.69 ±0.09 and 0.87 ±0.08, respectively. The sum of all POP congeners by clutches varied from 27.41 to 441.51 ng·g^−1^ lipid, and the average of all clutches was 121.86 ±98.5451 ng·g^−1^ lipid.

**Table 1.**
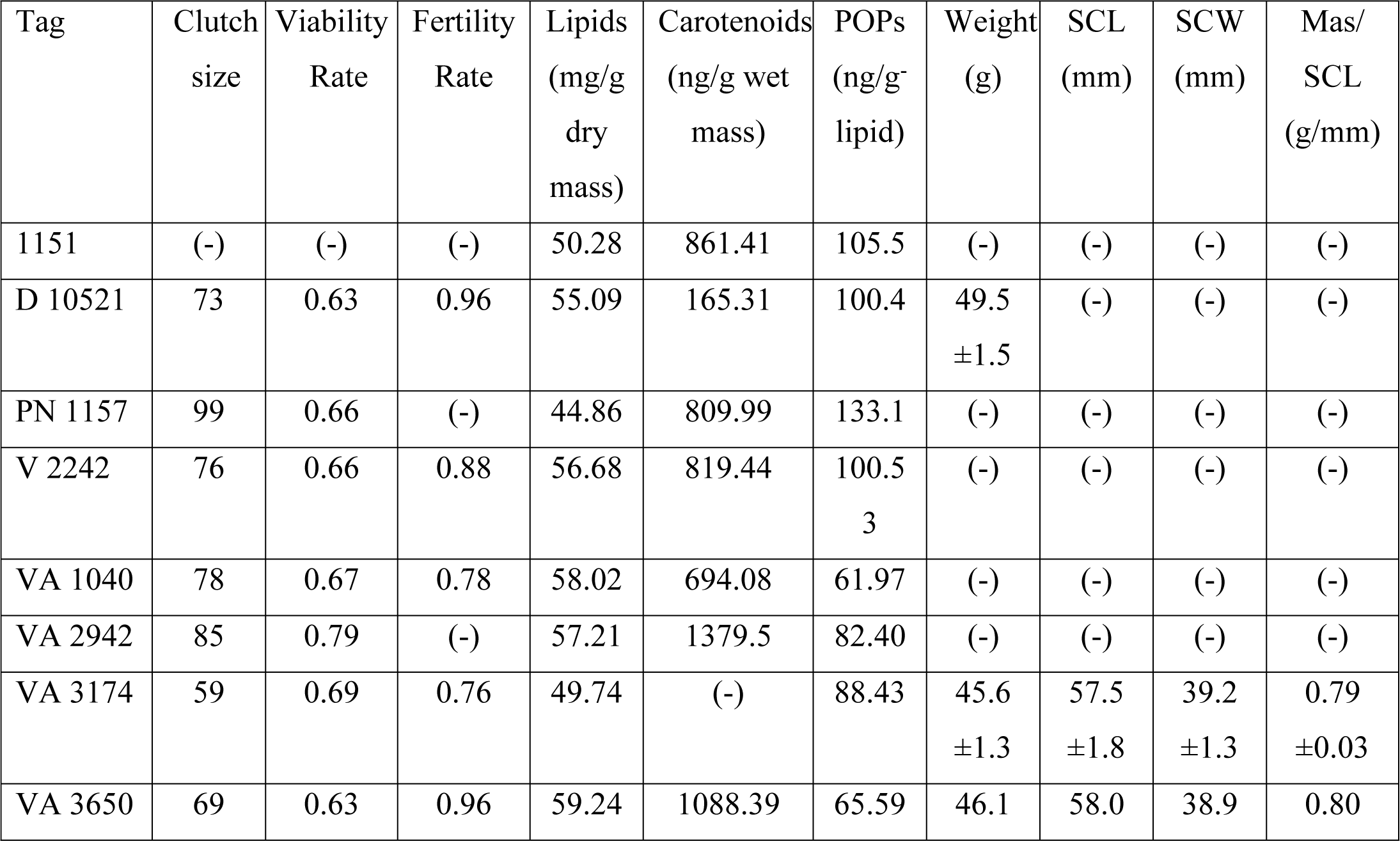
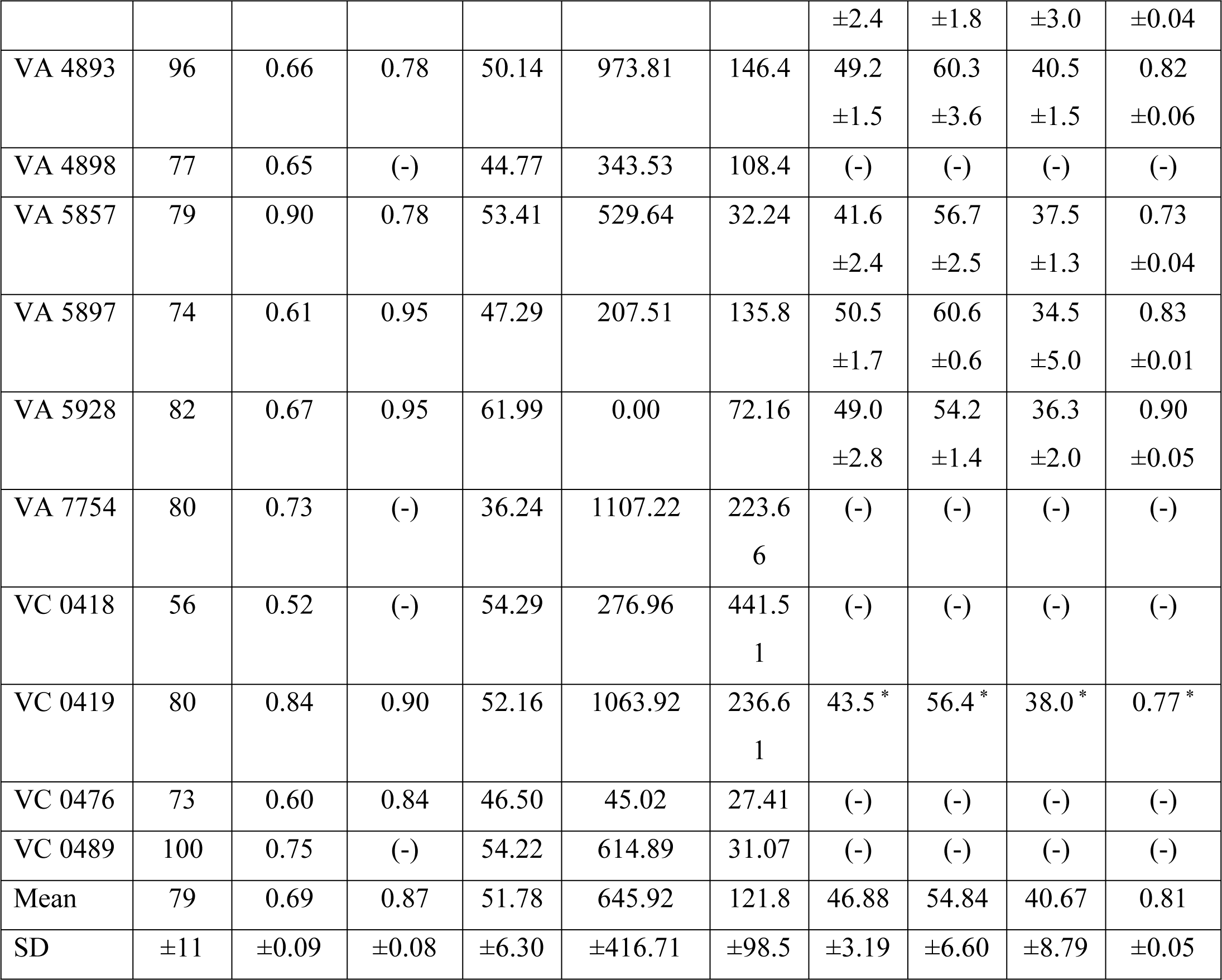
Reproductive parameters, egg yolk composition and hatchling morphometrics of leatherback turtles. Clutch size is equal to number of yolked eggs; viability rate calculated as clutch size divided by clutch size plus SAGs; fertility rate calculated as the proportion of fertile eggs. Total content of lipids, carotenoids and persistent organic pollutants (POPs) in egg yolks. Hatchling morphometrics data are mean ± SD of four to ten hatchlings per nest. SCL and SCW are straight carapace length and straight carapace width, respectively.

| Tag | Clutch size | Viability Rate | Fertility Rate | Lipids (mg/g dry mass) | Carotenoids (ng/g wet mass) | POPs (ng/g<sup>-</sup> lipid) | Weight (g) | SCL (mm) | SCW (mm) | Mas/ SCL (g/mm) |
| --- | --- | --- | --- | --- | --- | --- | --- | --- | --- | --- |
| 1151 | (-) | (-) | (-) | 50.28 | 861.41 | 105.5 | (-) | (-) | (-) | (-) |
| D 10521 | 73 | 0.63 | 0.96 | 55.09 | 165.31 | 100.4 | 49.5<br>$\pm 1.5$ | (-) | (-) | (-) |
| PN 1157 | 99 | 0.66 | (-) | 44.86 | 809.99 | 133.1 | (-) | (-) | (-) | (-) |
| V 2242 | 76 | 0.66 | 0.88 | 56.68 | 819.44 | 100.5<br>3 | (-) | (-) | (-) | (-) |
| VA 1040 | 78 | 0.67 | 0.78 | 58.02 | 694.08 | 61.97 | (-) | (-) | (-) | (-) |
| VA 2942 | 85 | 0.79 | (-) | 57.21 | 1379.5 | 82.40 | (-) | (-) | (-) | (-) |
| VA 3174 | 59 | 0.69 | 0.76 | 49.74 | (-) | 88.43 | 45.6<br>$\pm 1.3$ | 57.5<br>$\pm 1.8$ | 39.2<br>$\pm 1.3$ | 0.79<br>$\pm 0.03$ |
| VA 3650 | 69 | 0.63 | 0.96 | 59.24 | 1088.39 | 65.59 | 46.1 | 58.0 | 38.9 | 0.80 |
|  |  |  |  |  |  |  | ±2.4 | ±1.8 | ±3.0 | ±0.04 |
| VA 4893 | 96 | 0.66 | 0.78 | 50.14 | 973.81 | 146.4 | 49.2<br>±1.5 | 60.3<br>±3.6 | 40.5<br>±1.5 | 0.82<br>±0.06 |
| VA 4898 | 77 | 0.65 | (-) | 44.77 | 343.53 | 108.4 | (-) | (-) | (-) | (-) |
| VA 5857 | 79 | 0.90 | 0.78 | 53.41 | 529.64 | 32.24 | 41.6<br>±2.4 | 56.7<br>±2.5 | 37.5<br>±1.3 | 0.73<br>±0.04 |
| VA 5897 | 74 | 0.61 | 0.95 | 47.29 | 207.51 | 135.8 | 50.5<br>±1.7 | 60.6<br>±0.6 | 34.5<br>±5.0 | 0.83<br>±0.01 |
| VA 5928 | 82 | 0.67 | 0.95 | 61.99 | 0.00 | 72.16 | 49.0<br>±2.8 | 54.2<br>±1.4 | 36.3<br>±2.0 | 0.90<br>±0.05 |
| VA 7754 | 80 | 0.73 | (-) | 36.24 | 1107.22 | 223.6<br>6 | (-) | (-) | (-) | (-) |
| VC 0418 | 56 | 0.52 | (-) | 54.29 | 276.96 | 441.5<br>1 | (-) | (-) | (-) | (-) |
| VC 0419 | 80 | 0.84 | 0.90 | 52.16 | 1063.92 | 236.6<br>1 | 43.5 * | 56.4 * | 38.0 * | 0.77 * |
| VC 0476 | 73 | 0.60 | 0.84 | 46.50 | 45.02 | 27.41 | (-) | (-) | (-) | (-) |
| VC 0489 | 100 | 0.75 | (-) | 54.22 | 614.89 | 31.07 | (-) | (-) | (-) | (-) |
| Mean | 79 | 0.69 | 0.87 | 51.78 | 645.92 | 121.8 | 46.88 | 54.84 | 40.67 | 0.81 |
| SD | ±11 | ±0.09 | ±0.08 | ±6.30 | ±416.71 | ±98.5 | ±3.19 | ±6.60 | ±8.79 | ±0.05 |

### Fat and carotenoid levels

Lipid content of the egg yolk was averaged by nest (18 clutches) and the total mean accounted for 51.78 ± 6.3 mg·g^−1^ dry weight. Total concentration of fatty acid methyl esters (FAME) was 321.59 mg·g^−1^ dry weight of the total lipid content in egg yolk. Fatty acid profiles are presented in Table 2.

**Table 2.**
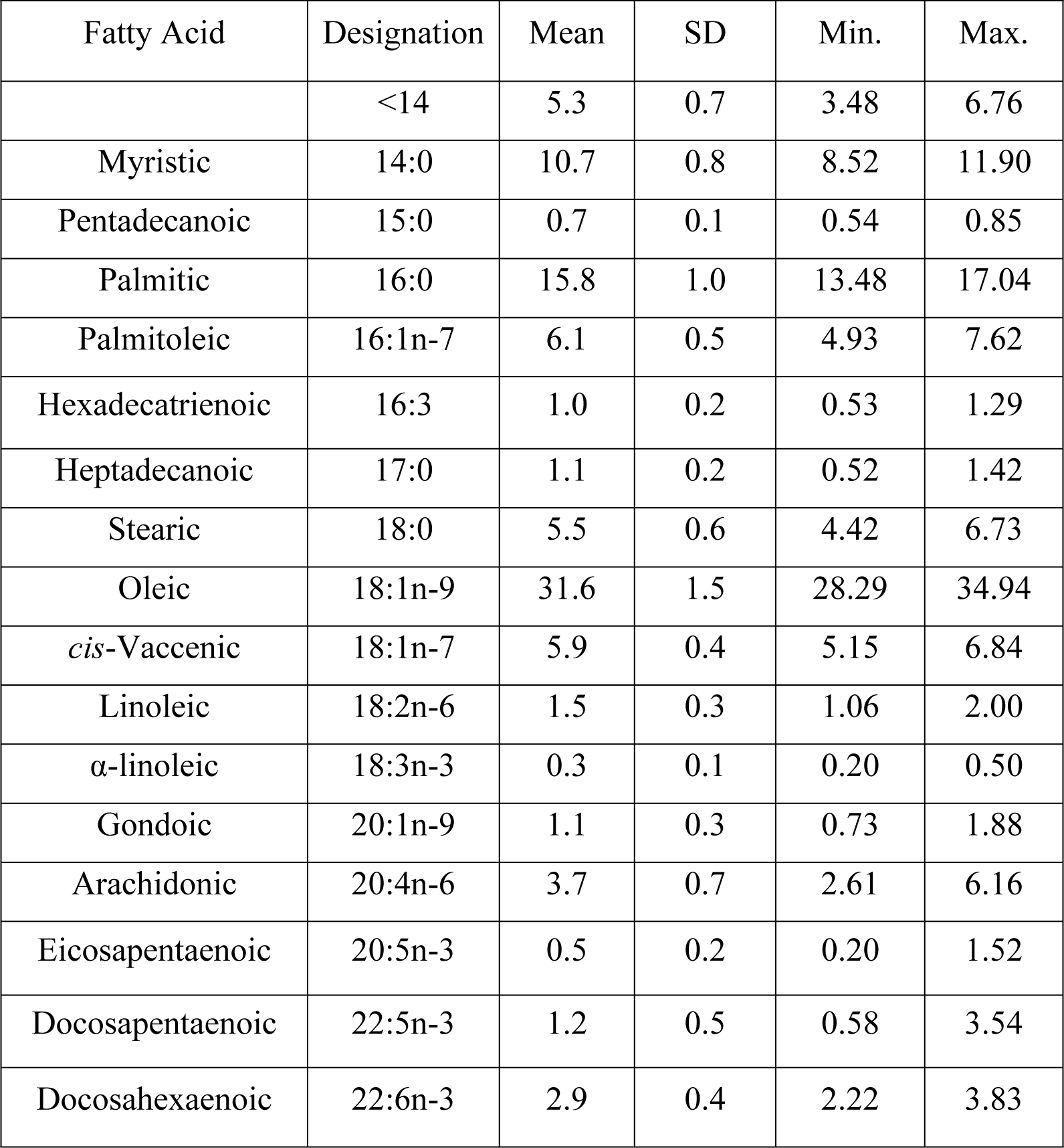
Fatty acid profiles of egg yolks from 18 clutches (N = 47 eggs) of leatherback turtles. Values are expressed as proportion of the total fatty acids (FAME) present in lipid content of egg yolks.

As usual in vertebrates, Oleic was the main fatty acid, representing approximately 30% of the total. Palmitic and Myristic had also an important percent, of about 16 and 11%, respectively. Palmitoleic, *cis*-Vaccenic and Stearic showed similar proportions around 6%. Arachidonic presented a relatively high proportion of 3.7% when compared to other turtle species. Docosapentaenoic and Docosahexaenoic fatty acids were also highly represented (1.2 and 2.9%, respectively) if contrasted with the proportions reported before for *Dermochelys coriacea* of those fatty acids. When fatty acids are grouped by class (Table 3), monounsaturated fatty acids accounted for almost half of the total fatty acids (45.1%), followed by saturated (33.8%). Polyunsaturated fatty acids represented around 12%, with the mean of n-3 being slightly lower than that of n-6 (5.1 *vs.* 5.6%).

**Table 3.**
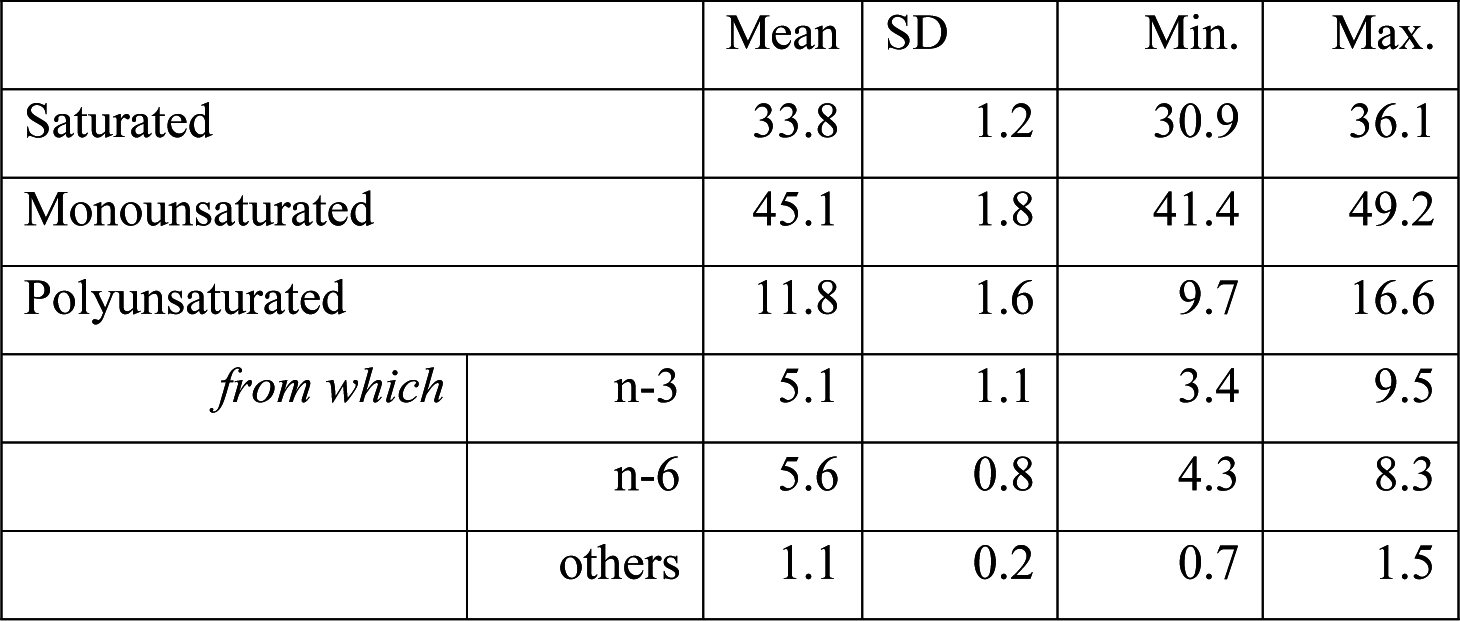
Fatty acid proportions of leatherback turtle eggs grouped by class, obtained from egg yolks of 18 clutches (*N = 47 eggs*).

Regarding to carotenoid levels in eggs, averaged concentrations ranked from 0.00 to 1107.22 ng·g^−1^ wet weight and mean value of 645.92 ±416.71 ng·g^−1^ wet weight. Zeaxanthine was the main of these components with mean values of 481.06 ng·g^−1^ wet weight and maximum values of 1220.30 ng·g^−1^ wet weight (Table 4). In second ranking order, an apparently unknown carotenoid termed unidentified 1 reached values up to 652.20 ng·g^−1^ wet weight and had mean concentration of 156.63 ng·g^−1^ wet weight. Both pigments had a high number of above-zero cases (38 and 34 eggs, respectively). On the other hand, Lutein was poorly represented in terms of both above-zero cases and mean concentrations (Table 4).

**Table 4.** Concentration (ng·g^−1^ wet mass) of carotenoids in egg yolks of leatherback turtles (N=47 eggs). n (C > 0) indicates the number of eggs that showed above-zero values of carotenoid concentrations.

| | $n$ ($C > 0$) | Mean | SD | Min. | Max. |
| --- | --- | --- | --- | --- | --- |
| Zeaxanthine | 38 | 481.06 | 342.2 | 0.00 | 1220.30 |
| Lutein | 8 | 32.92 | 79.72 | 0.00 | 358.57 |
| Unidentified 1 | 34 | 156.63 | 152.18 | 0.00 | 652.20 |
| Unidentified others | 11 | 26.12 | 49.80 | 0.00 | 192.61 |
| Total Carotenoids | 41 | 696.73 | 488.51 | 0.00 | 1811.68 |

### Variability

Lipid and carotenoid profiles in the present study were not significantly different between clutches (*p* > 0.05). This could be due to the large deviation also presented within some clutches. ANOSIM-R values were 0.47 for lipids and 0.31 for carotenoids, with all *p-values* > 0.05. NMDS plots better illustrated the variation-grouping pattern within and among clutches (Fig. 1). Note that it does exist grouping in eggs of some clutches (e.g. clutches 1, 2, 4 and 8 for lipids) (Fig. 1a). Carotenoid levels slightly differed both within and among clutches and that is the reason for almost all eggs plotting together (Fig.1b).

**Figure 1.**
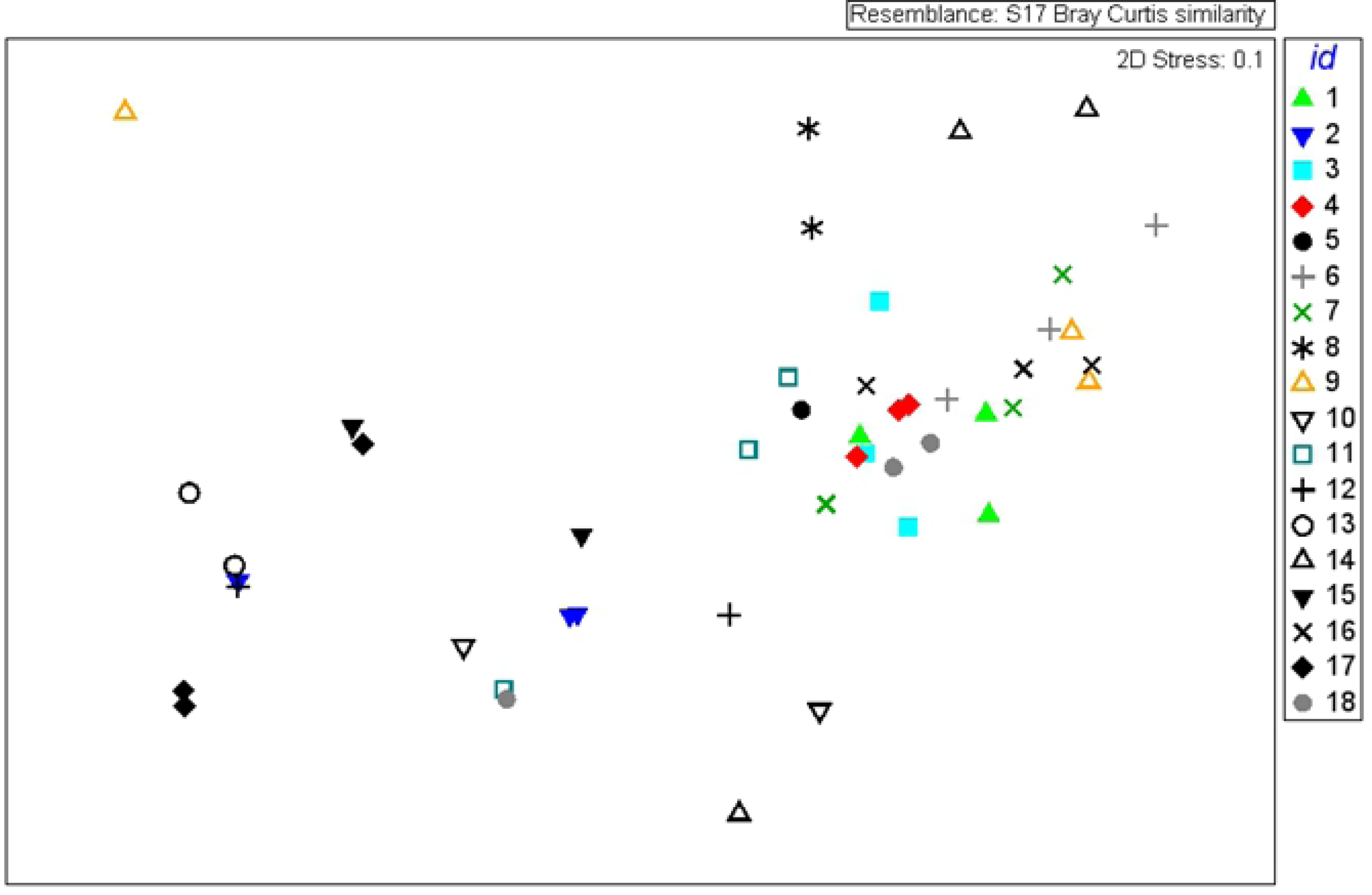
Dermochelys coriacea. 2D-NMDS plots of egg concentration profiles of: a) lipid and fatty acids; b) carotenoids. Eggs of the same clutch are indicated by the same symbol, and each clutch has an assigned id. Non-grouping of eggs from the same clutch indicates that variation within clutches is larger than variation among clutches. Stress = 0.1 suggests good fit; Stress = 0, poor fit.

Lipid content, each class of fatty acids, and zeaxanthine were averaged by nest and considered as a variable. Mean nest values of the different POP congeners were taken from De Andrés et al [36]. Shapiro-Wilk (S-W) tests for all variables indicated that most of them were normally distributed (*p* > 0.05) except for some POP groups (hepta- to deca-BDEs and all PCB groups). Therefore, concentrations of those POP groups were log10-transformed and S-W normality test performed. If the obtained *p-values* were > 0.05, parametric correlations were then executed between all different variables. Significant and positive relationship was only found between polyunsaturated fatty acids (PUFA) (stronger for the n-6 PUFA) and lipid-normalized POP concentrations (*p* < 0.05) (Fig. 2).

**Figure 2.**
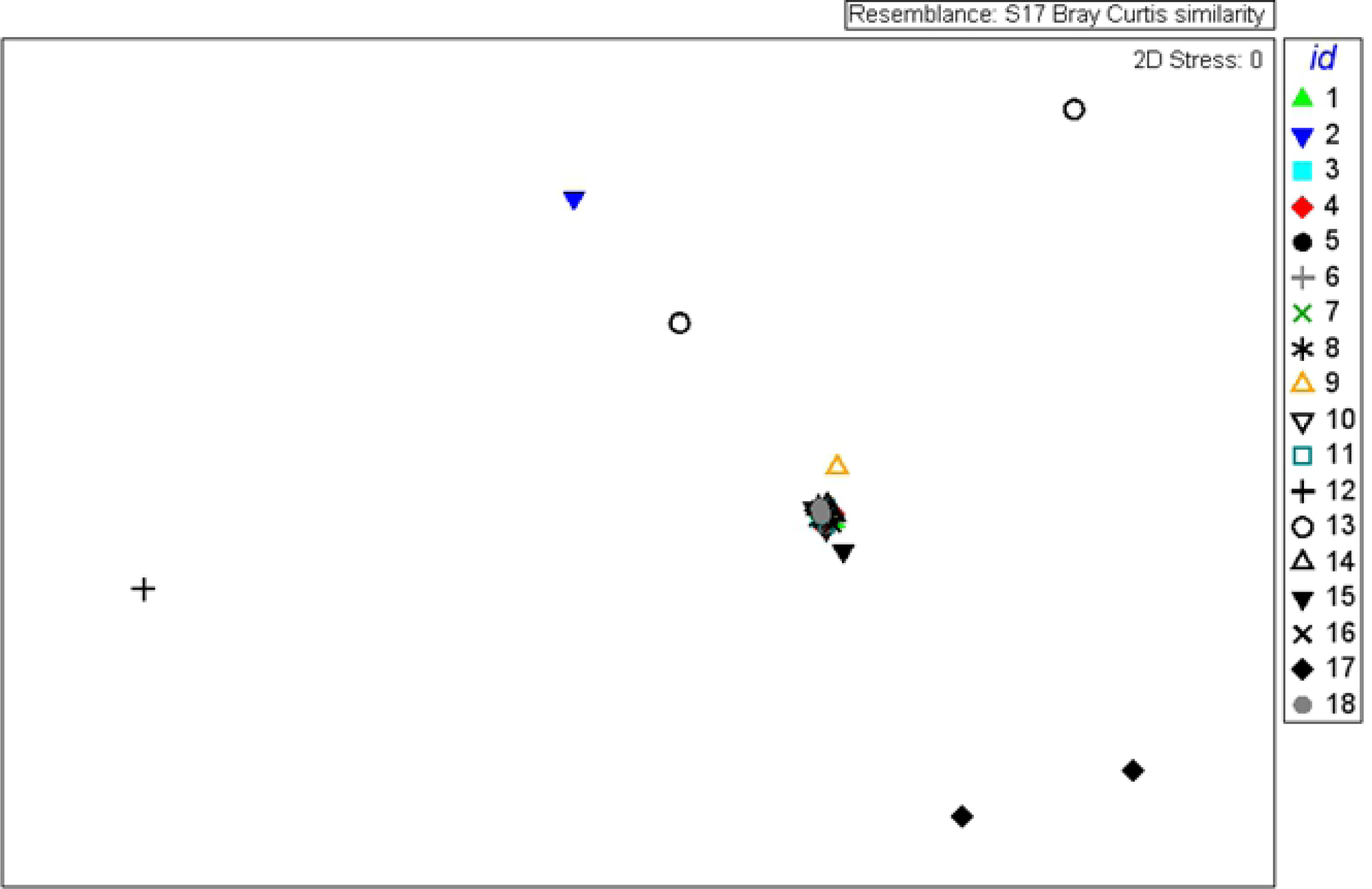
Significant relationship between POP concentrations and polyunsaturated fatty acids (PUFA) in eggs of leatherback turtles, from which b) n-6 series were also significantly related to POP concentrations. Values were averaged by nest (mean ± SD). Pearson-R coefficient, p-significance and linear trendline.

Reproductive parameters and hatchling morphometrics were all normally distributed (S-W, *p* > 0.05), except for the fertility rate. Thus, all correlations related to the fertility rate were non-parametric (Spearman rank order). S-W normality test was also realized for all variables taking into account the nest ranks covered by both variables; namely, hatchling parameters were only available for 7 to 8 nests, so S-W test was executed for all the other variables within those nests in order to choose the best correlation method. And so on for all possible combinations.

CCL of nesting females was positive correlated to saturated fatty acids (R = 0.51, *p* < 0.05), indicating that larger females may accumulate more of these fatty acids. Regarding to reproductive parameters two biologically-relevant and significant correlations were found (Fig. 3). Firstly, obtained from Pearson-parametric test, zeaxanthine concentrations correlated positively to the viability rate (R = 0.52; *p* < 0.05). Secondly, obtained from non-parametric test (Spearman-rank order), PUFA n-3 well correlated to the fertility rate (R = 0.75; *p* = 0.008). These findings are further discussed below.

**Figure 3.**
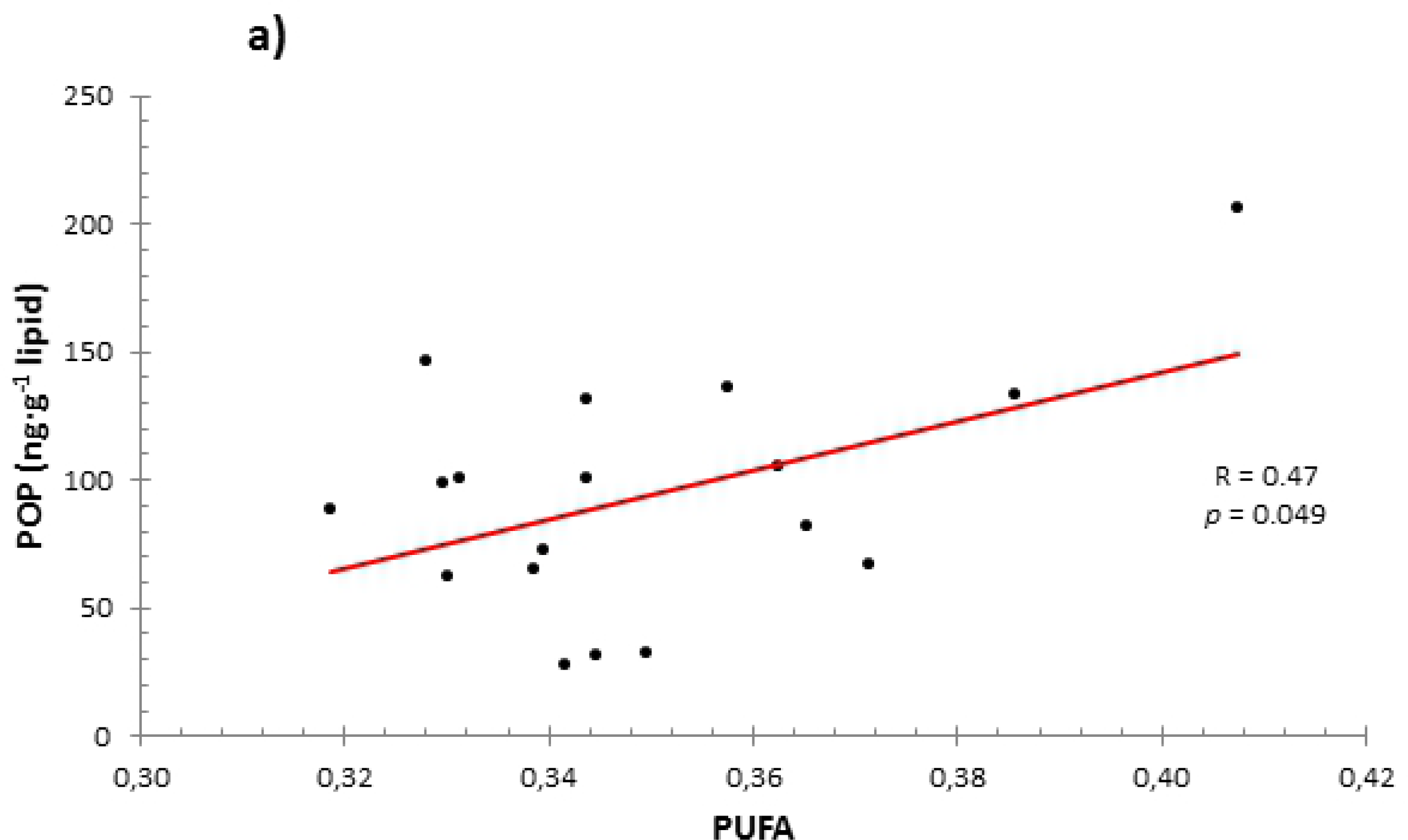
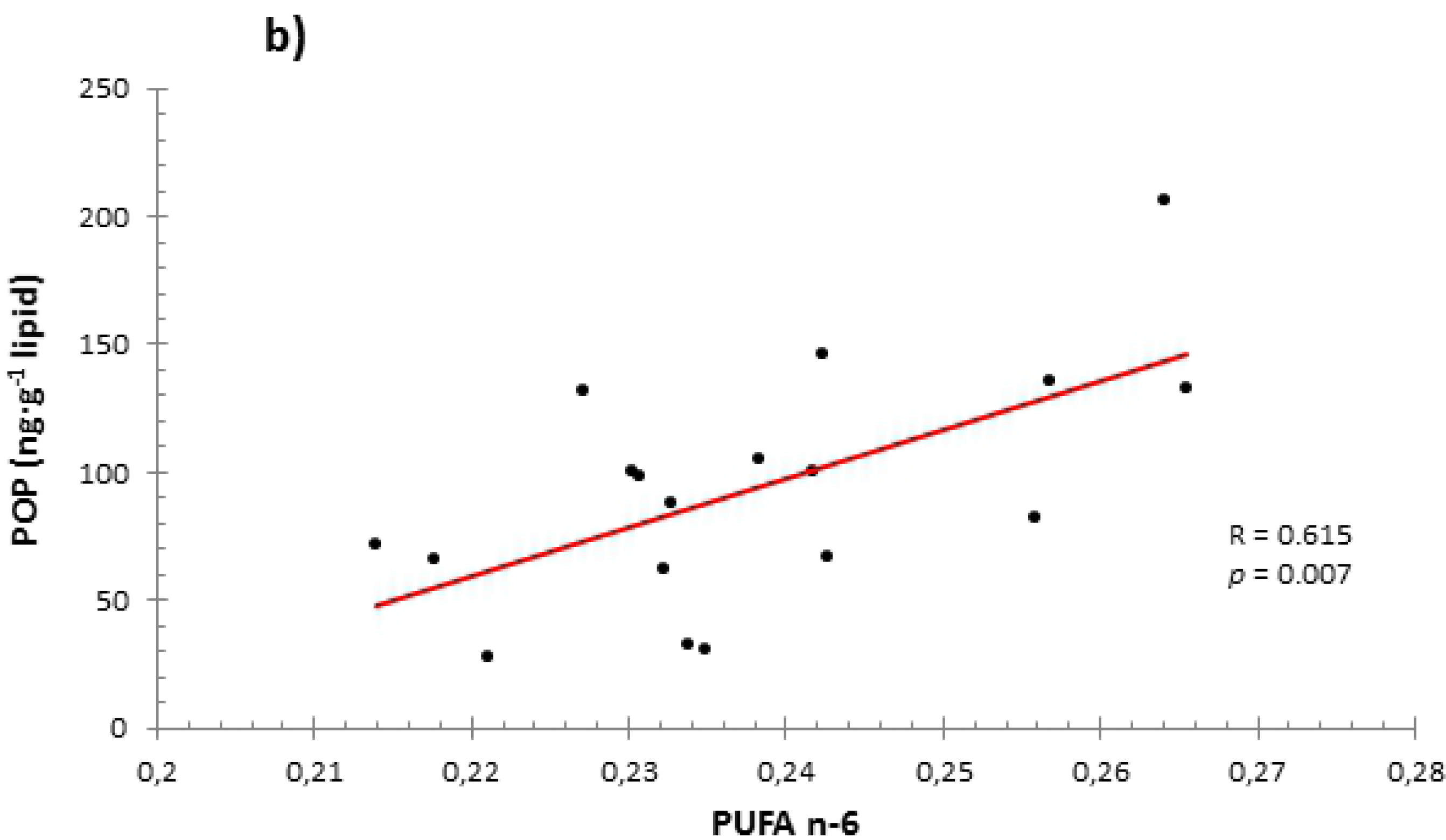
Relationships between different compounds averaged by nest (mean ± SD) and reproductive parameters of leatherback turtles. a) Parametric correlation of Zeaxanthine concentrations and Arcsin-transformed viability rate (N = 16); b) Non-parametric Spearman-rank correlation of PUFA n-3 percentage against Fertility rate (N = 11). Note that both fertility and n-3 are proportions, so none of them had been transformed.

No significant relationships were found between either carotenoids or POPs and hatchling morphometrics. However, fatty acid classes interacted with hatchling parameters in different ways. For example, the higher proportions of n-6 fatty acids, the larger SCL of hatchlings (R = 0.83; *p* < 0.05). Similarly, high proportions of saturated fatty acids, resulted in larger sizes regarding to SCW (R = 0.92; *p* = 0.003) (Fig. 4). CCL of nesting female was also positively related to hatchling SCW, suggesting that the larger the mother, the wider the offspring (R = 0.82; *p* < 0.05) (Fig. 5). Moreover, it was found that hatchling mass (weight) was strongly and negatively correlated to the viability rate (R = −0.896; *p* = 0.003), and the mass:SCL ratio to the hatching rate (R = −0.94; *p* = 0.002), indicating a potential strategy of reproduction success (Fig. 6).

**Figure 4.**
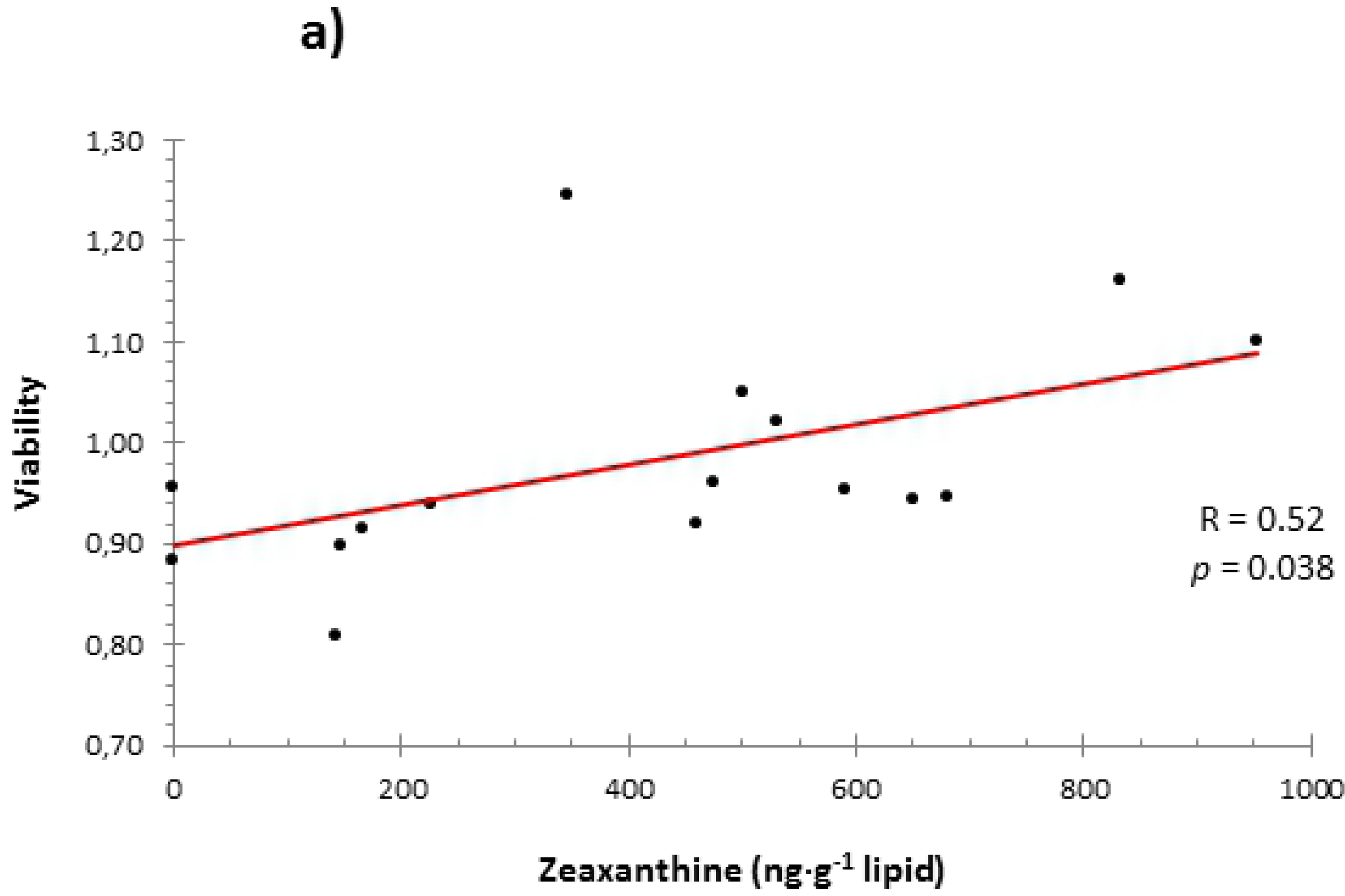
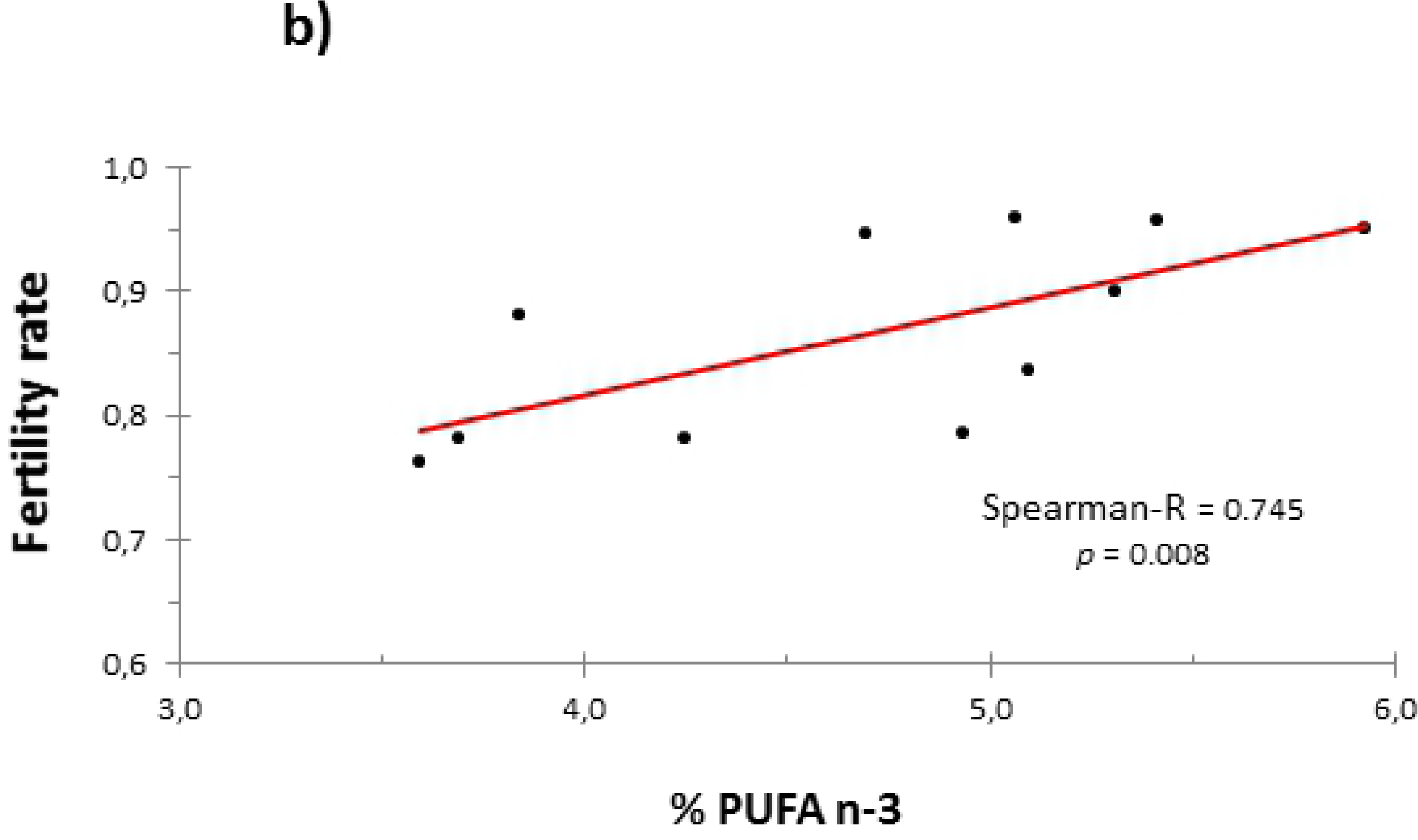
Relationship between fatty acids and hatchling size of leatherback turtles. a) Arcsin-transformed PUFA n-6 proportions against straight carapace length (SCL); b) Arcsin-transformed saturated fatty acid proportions against straight carapace width (SCW).

**Figure 5.**
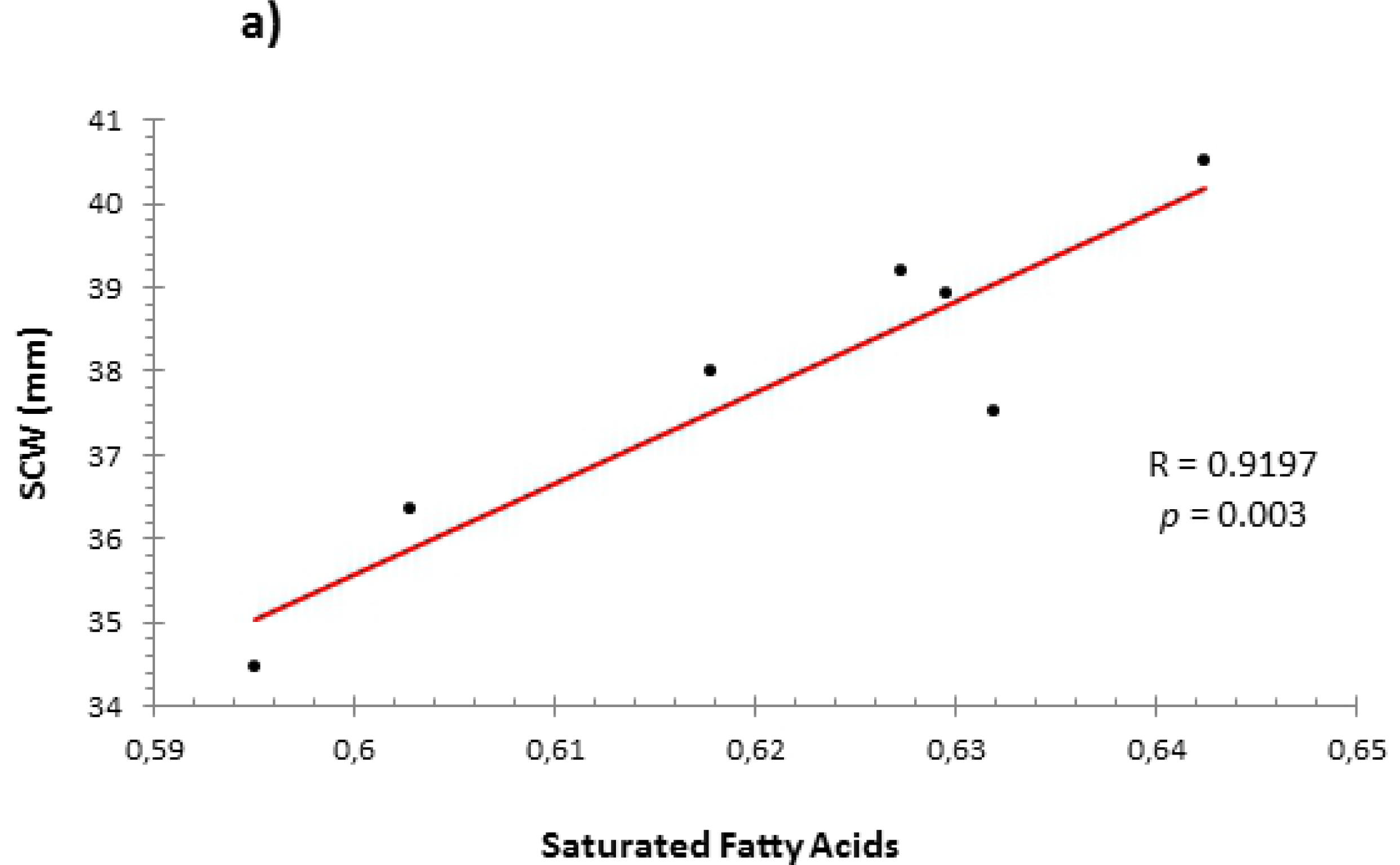
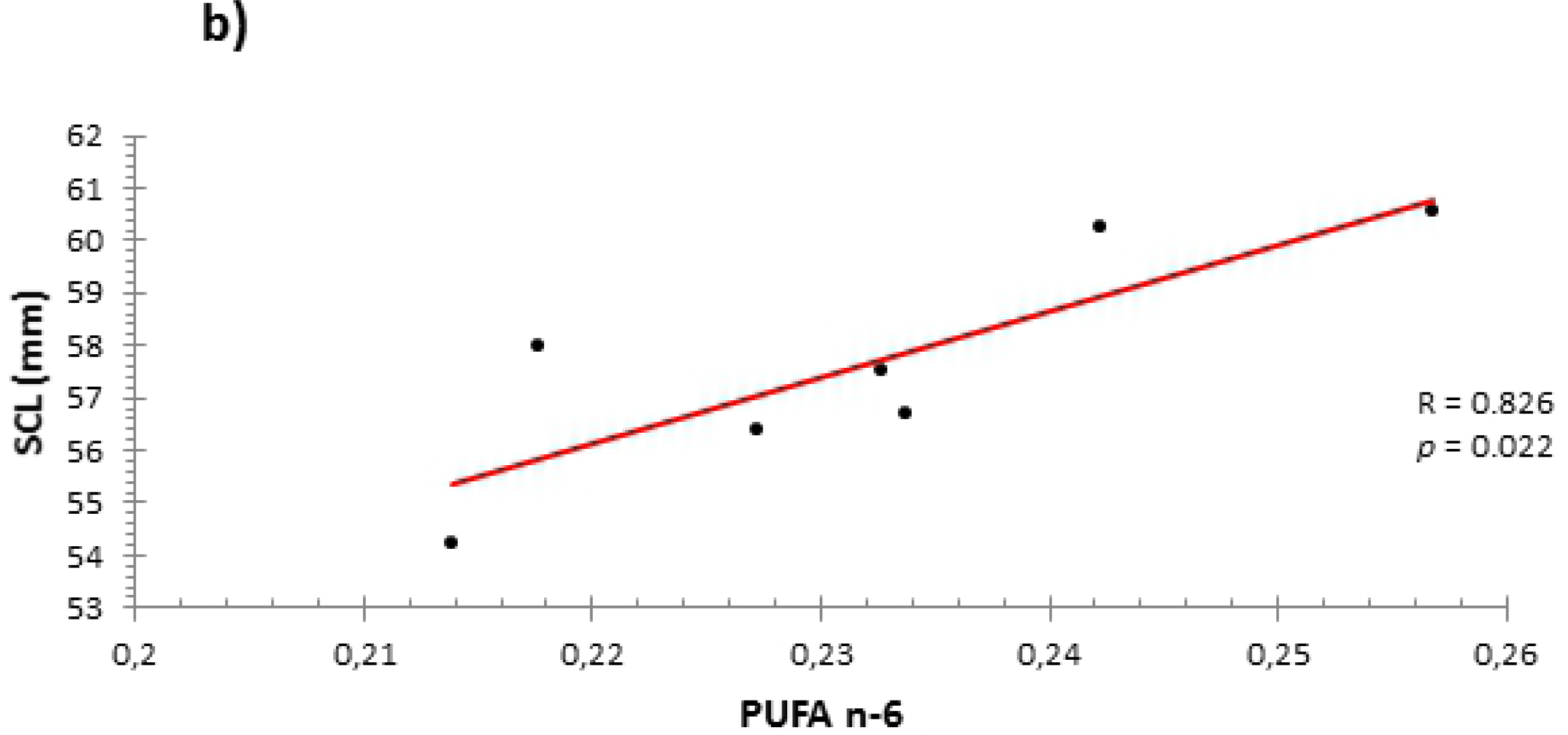
Relationship between nesting female and hatchling sizes of leatherback turtles. CCL = curved carapace length; SCW = straight carapace width. Pearson-R coefficient, *p*-values and linear trend line equation are shown.

**Figure 6.**
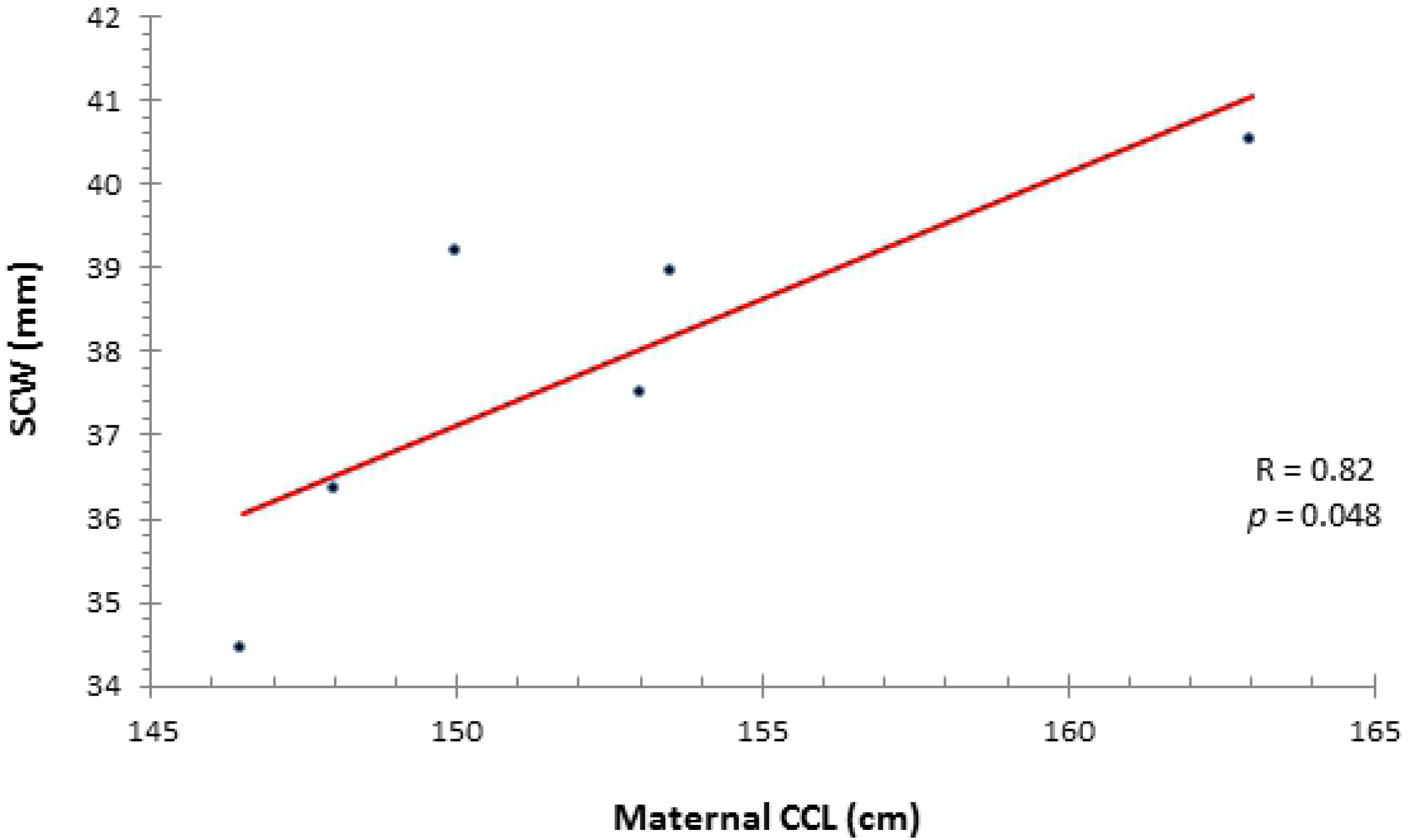
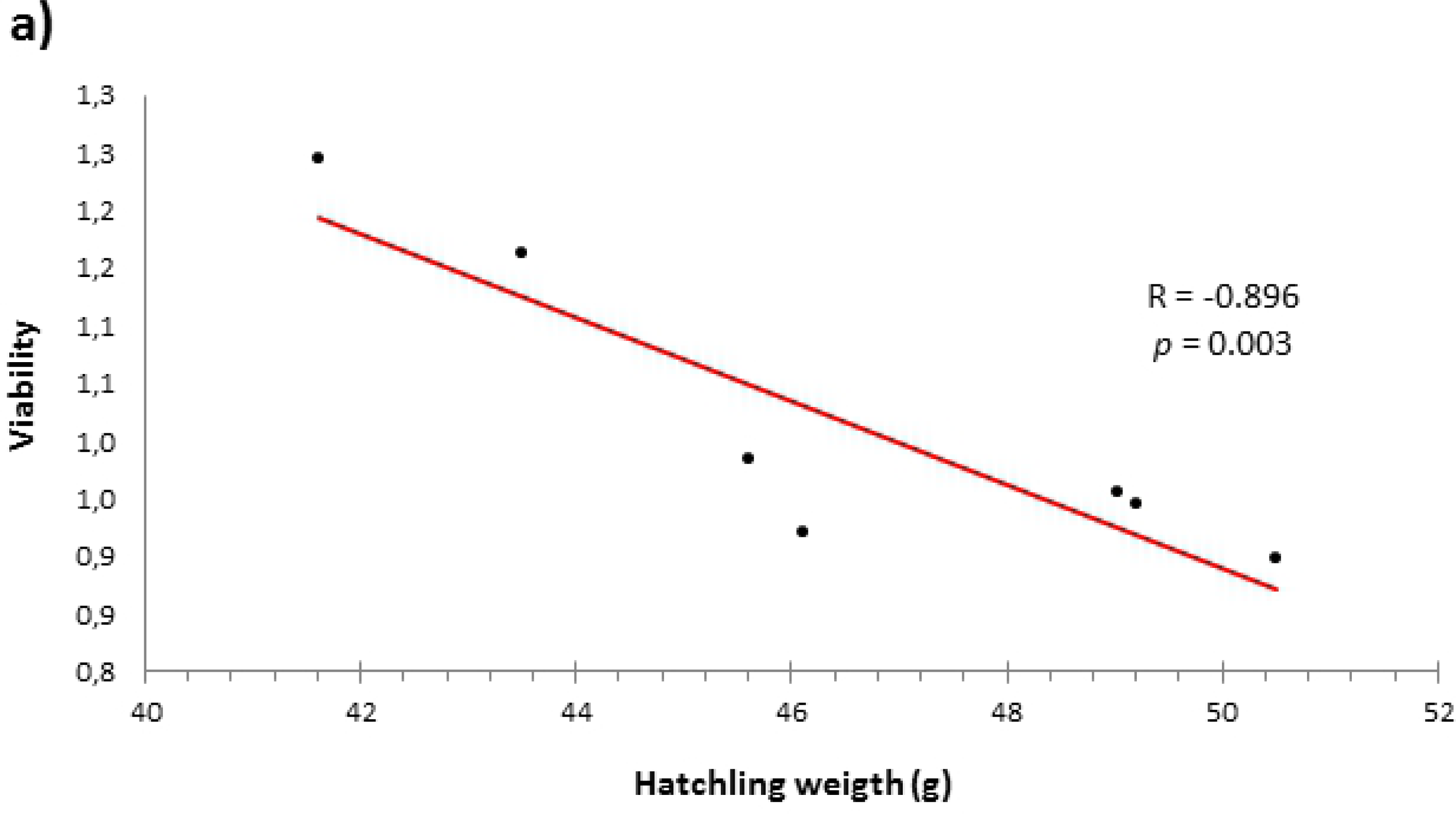
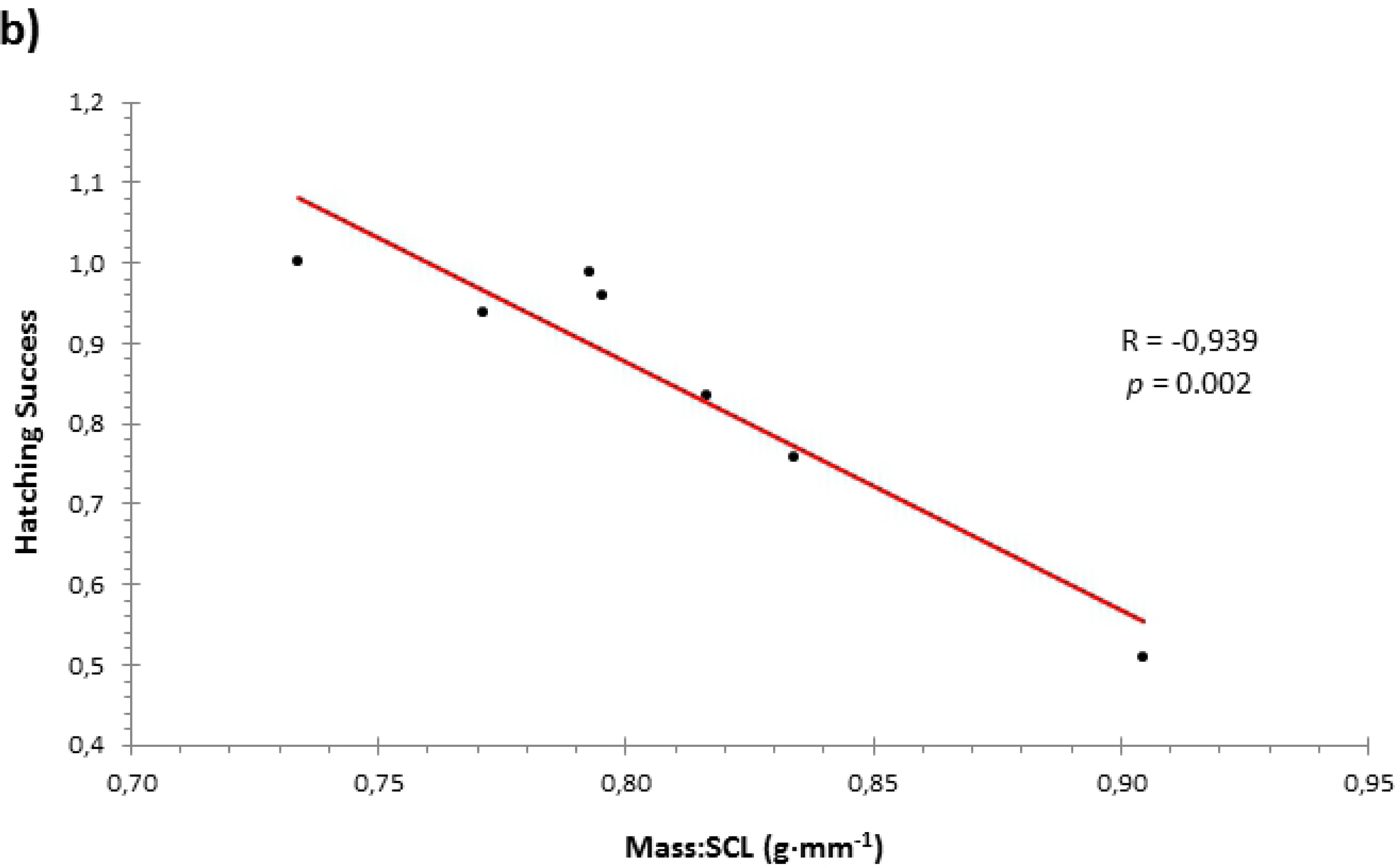
Relationships between hatchling morphometrics and reproductive parameters of leatherback turtles. a) Correlation of Arcsin-transformed viability rate against hatchling mass (weight); b) correlation of Arcsin-transformed hatch rate against mass:LRC ratio. Pearson-R, *p* values and linear fit equations are shown.

## DISCUSSION

### Variability within nests

According to the results of ANOSIM, it was demonstrated that for the analysis of all lipids and fatty acids the sample size per nest was insufficient. That is, the internal variability of some nests was higher compared to the values found in other nests. Still, there were nests that showed a remarkable degree of clustering, as shown in the NMDS plots. Possibly, there are some factors influencing this variability, such as the quality of the foraging areas, travelled distances, temperature of waters, type of available food, maturity of the laying females, etc. In the case of carotenoids, the concentrations are low, and the intra-nest and inter-nest variabilities are of the same order and both agglutinate in a small region. Such low values are likely the result of a poor-pigmented food diet, which consist in jellyfish and gelatinous phytoplankton [22, 23, 24].

### Importance of fatty acids in reproduction

Fats are essential for reproduction in turtles. In this study, we observed a strong positive and significant relationship between saturated fatty acids and curved carapace length of nesting females. Given that many of stored fat reserves (which will be used subsequently in the production of eggs) are at sub-carapace level [37] and that the saturated represented about 35% of total fatty acids, such correlation between stem length and amount of saturated fatty seems to be quite meaningful.

Around 80% of the fat comprising the liver serve as a source of energy available for the production of eggs. The mechanism of mobilization of fats for reproductive functions begins with the conversion of such fat stores in the liver to start vitellogenesis and for ovarian follicles to get rich in lipids and fatty acids, which will make up much of egg yolk [45]. Therefore, the relationship found in this study between fertility and proportion of n-3 fatty acids appears consistent. Similarly, a positive relationship between fertility and the proportion of eicosapentaenoic acid (20:5 n-3) was found by other authors [46] in prawns. Studies in green turtle (*C. mydas*) have shown that lipids account for almost 70% of the egg yolk [45], since this is the main source of energy used for embryo development and also a major energy source for hatchlings to develop their first activities towards reaching safer foraging areas in open waters [47]. In addition, fatty acids are essential in the construction of new cells as well as in supplying the energy demand of the embryo. Therefore, these fatty acids seem to be essential for the proper development of the embryonic tissues, assuming a relevant factor in the offspring success [48]. Although it has been suggested that small deficiencies in n-3 polyunsaturated fatty acids (PUFAs) may be the cause of low rates of hatching in oysters [49], birds [50, 51] and alligators [44, 48], these relationships were not found for leatherbacks in this study. However, it has been obtained here a strong positive relationship between n-6 PUFAs and length of hatchlings. This could be due to the relatively high levels of arachidonic acid (20:4 n-6) present in the eggs of *D. coriacea* analyzed here (3.7% ± 0.7%), which were around two to three times the levels obtained in eggs of *C. mydas* (1.19%)[45]; *T. hermanni* (1.5%) [27]; or *E. maquarii* (1.8%) [26]; Thompson *et al*., 1999). The arachidonic acid is found in the cell membranes of embryonic organs such as the brain, liver and heart, and it is believed to play a regulatory function [50, 53]. Therefore, the results obtained here complement the information about the importance of arachidonic acid in the growth of the embryo, suggesting that high levels provide larger hatchlings, conferring them a greater chance of survival [54]. The positive relationship between the proportion of saturated fatty acids and carapace width of hatchlings would be justified by the length of the mother (which also has a positive relationship with these unsaturated acids, as suggested above), since a positive relationship between stem length and hatchling width was obtained. It seems that POPs have some affinity with PUFAs, since they were positively and significantly correlated. PUFAs are characterized by having more than one double bond. Considering that POPs have a strong lipophilic character and that PUFAs have a greater tendency to bind to other molecules, it is plausible that this positive relationship actually happen. The harmful effects that POPs may have in several aspects of reproduction success have been widely reported in sea turtles [31, 32, 33, 34], and specifically in *D. coriacea* [35, 36]. Therefore, it must be borne in mind that the existence of a positive relationship between POPs and PUFA (especially n-6) could probably cover up the negative effects of POPs, since both PUFAs n-3 and n-6, as previously argued, have positive effects on reproductive success. Further studies of these interactions and their potential effects on reproduction of sea turtles are required.

### Importance of carotenoids in embryonic development and viability

Carotenoids are lipid-soluble antioxidants which are thought to protect the embryonic tissues from peroxidative damage [55]. Despite the importance that carotenoids have in embryonic development, there is not much information on sea turtle species. Although there have been many studies on carotenoid concentration in domestic chicken eggs, it is only during the past decade that similar information has begun to accumulate for reptile eggs [26, 27, 53] and only one of them refers to sea turtle eggs, namely *Chelonia myda*s [28]; in which it was demonstrated that the concentration of carotenoids may influence the quality of the offspring phenotype (the higher the carotenoid concentrations, the larger the hatchlings, in terms of SCL). In our study, zeaxanthine was presented as the most abundant pigment, with concentrations of 0.48 ± 0.34 µg·g^−1^ fresh yolk and total carotenoid concentrations of 0.7 ± µg·g^−1^ fresh yolk. Such concentrations are around 10 times lower than those reported in green turtles by Weber [28](2010) (5.14 ± 0.4 µg·g^−1^ fresh yolk), in samples with very similar zeaxanthine and lutein proportions as apposite to our study. Moreover, the total concentrations of carotenoids reported by Thompson *et al*. [26] and Speake *et al*. (2001) for tortoise and freshwater turtles (*Testudo hermanni* and *Emydura maquarii*, respectively) were significantly higher, reaching values of 9.8 and 12.6 µg·g^−1^ fresh yolk, respectively. In *T. hermanni*, lutein was the main pigment, with concentrations of 8.7 µg·g^−1^ fresh yolk. These comparative results suggest that *Dermochelys coriacea* feeds on low-pigments diets, which would be supported by the fact that they basically consist on jellyfish, and these do not apparently have large amounts of these pigments. In this study we observed that zeaxanthine had a significant and positive relationship with the viability rate. Due to the association of low number of yolked eggs, the low concentrations of carotenoids found in eggs of this species and the importance of carotenoids for embryonic development in the leatherback turtle, it is feasible to think that this relationship may indicate that zeaxanthine might be acting as a limiting factor during vitellogenesis in this species. This would ensure some minimum requirements for the formation of the egg yolk and the proper development of the embryo. Furthermore, the high variability found in carotenoid concentrations among different nests suggest that females were in unequal physiological conditions. This would be supported by the evidence found by Weber [28], where carotenoid concentrations significantly decrease across successive clutches laid by an individual female within a nesting season. Like for fatty acids, more research is needed to clarify and ascertain whether carotenoids may somehow play a limiting role on vitellogenic processes in leatherback turtles.

## CONCLUSIONS

Studies of reproductive success and its potential influencing factors are strictly necessary in order to know the population trend of this critically endangered species (*Dermochelys coriacea*).

This study provides an important baseline of lipid content, fatty acid profiles, and carotenoid concentrations, which are all considered as fundamental in reproduction physiology. They will serve as baseline data for future studies in this field.

Results from this study also provide strong evidence that all fatty acid proportions and carotenoid concentrations may be influencing the reproductive success in leatherback turtles, in different ways. Thus, it appears that n-3 PUFAs contribute to embryonic formation at the most initial stages, increasing the fertility rate of nests. Furthermore, higher proportions of n-6 PUFAs may favor the successful development of the embryo, conferring a larger length to hatchlings and thereby increasing their potential survival. Moreover, the concentrations of zeaxanthine in leatherback turtle eggs were found to be extremely low, likely due to their diet based on jellyfish, which is poor in pigments. This pigment is essential as an antioxidant in embryonic development and, apparently, may act as a limiting factor in the process of vitellogenesis, since a positive correlation was found between concentrations of zeaxanthine and viability rate.

We also found a strong positive relationship between PUFA and POPs, suggesting a higher affinity of contaminants to such polyunsaturated fatty acids. Given that both types of molecules have shown antagonistic effects in reproductive parameters; this relationship could cover part of the harmful effects which POPs may have in reproduction.

These facts show the complexity inherent in this type of studies on assessing the possible influence of compounds (of different nature and with different properties) in reproduction. We recommend that a global overview must be present and that more integrated and comprehensive studies must be undertaken. Further investigations will be necessary considering different physiological levels that reproduction system may be implicated on, and various potential interactions between these lipophilic compounds, in order to obtain more accurate and conclusive results.

## AKNOWLEDGEMENTS

We thank all the volunteers from Pacuare Reserve, Didier Chacón and Hilda Denham; Zaida Fernández and Carlos Hernández from Endangered Wildlife Trust. Financial support came from the BBVA fundation, Consejo Superior de Investigaciones Científicas of Spain (CSIC) and the Universidad Internacional Menéndez Pelayo (Madrid, Spain).

